# Evaluation of the EUROIMMUN Anti-SARS-CoV-2 ELISA Assay for detection of IgA and IgG antibodies

**DOI:** 10.1101/2020.05.11.089862

**Authors:** Scott M. Matushek, Kathleen G. Beavis, Ana Abeleda, Cindy Bethel, Carlissa Hunt, Stephanie Gillen, Angelica Moran, Vera Tesic

## Abstract

As the Coronavirus 2019 (COVID-19) pandemic evolves, the development of immunoassays to help determine exposure and potentially predict immunity has become a pressing priority. In this report we present the performance of the EUROIMMUN enzyme-linked immunosorbent assay (ELISA) for semi-quantitative detection of IgA and IgG antibodies in serum and plasma samples using recombinant S1 domain of the SARS-CoV-2 spike protein as antigen. Specimens from patients, with and without COVID-19 infection, were tested at the University of Chicago Clinical Microbiology and Immunology Laboratory. Of 57 samples from COVID-19 PCR-negative patients, including 28 samples positive for common human coronavirus strains, 53 tested negative and 4 tested positive for IgA (93.0% agreement) while 56 tested negative and 1 tested positive for IgG (98.2% agreement). For COVID-19 PCR-positive patients, 29 of 30 (96.7%) samples collected ≥3 days after positive PCR were positive for IgA, and 28 of 28 samples collected ≥4 days after positive PCR were positive for IgG.

The EUROIMMUN Anti-SARS-CoV-2 ELISA Assay demonstrates excellent sensitivity for detection of IgA and IgG antibodies from samples collected ≥3 days and ≥4 days, respectively, after COVID-19 diagnosis by PCR. This assay did not demonstrate cross reaction in any of the 28 samples from patients with common human coronaviruses, including types HKU1, NL63, CV229E, and OC43.

## Introduction

In December 2019 a novel coronavirus emerged as the cause of severe respiratory disease and quickly spread causing a worldwide pandemic. Severe Acute Respiratory Coronavirus 2 (SARS-CoV-2) was determined to be the agent of coronavirus disease 2019 (COVID-19). The virus belongs to the Betacoronavirus genus of the Coronaviridae family, which also includes Severe Acute Respiratory Syndrome Coronavirus 1 (SARS-CoV-1) and Middle East respiratory Syndrome Coronavirus (MERS-CoV) (1). For diagnostic purposes many nucleic acid amplification assays were quickly developed and received Emergency Use Authorization (EUA) from the US Food and Drug Administration (FDA). Multiple manufacturers are offering serological assays, but few have received Emergency Use Authorization from the Food and Drug Administration (FDA); the EUROIMMUN IgG assay has received Emergency Use Authorization. Serological testing may be useful in conjunction with other laboratory tests and clinical findings of COVID-19 infection for epidemiological monitoring and outbreak control. Of the immunoassays currently available, choice of SARS-CoV-2 target antigens include the spike protein (S) or the nucelocapsid (N) (2). IgA antibodies can show higher sensitivity, while IgG antibodies typically have longer duration, better specificity, and are better suited for serosurveillance studies (3-5).

## Materials and Methods

The EUROIMMUN Anti-SARS-CoV-2 assay is an enzyme-linked immunosorbent assay (ELISA) that provides semi-quantitative in vitro determination of human antibodies of immunoglobulin classes IgA and IgG against severe acute respiratory syndrome coronavirus 2 (SARS-CoV-2) in serum or EDTA plasma (6-7).

Each kit contains microplate strips with 8 break-off reagent wells coated with recombinant structural protein of SARS-CoV-2. In the first reaction step, diluted patient samples are incubated in the wells. In the case of positive samples, specific antibodies will bind to the antigens. To detect the bound antibodies, a second incubation is carried out using an enzyme-labelled antihuman IgA or IgG (enzyme conjugate) catalyzing a color reaction. Results are evaluated semi-quantitatively by calculation of a ratio of the extinction of the control or patient sample over the extinction of the calibrator. This ratio is interpreted as follows:

- Ratio < 0.8 negative
- Ratio ≥ 0.8 to <1.0 borderline
- Ratio ≥ 1.1 positive

The IgG assay has received Emergency Use Authorization from the US Food and Drug Administration (FDA).

The University of Chicago Medicine uses 2 different RT-PCR assays allowed by the FDA under Emergency Use Authorization. The Roche cobas 6800 SARS-CoV-2 assay relies on amplification of the SARS-CoV-2 specific ORF1 gene as well as a portion of the E-gene conserved across the sarbecoviruses, a subgenus of coronaviruses which includes SARS-CoV-2. The Cepheid Xpert Xpress SARS-CoV-2 assay also detects the pan-sarbecovirus E-gene but uses the SARS-CoV-2 specific N-gene rather than ORF1 as its primary target. Samples tested include nasopharyngeal and nasal mid-turbinate swabs transported in viral transport or liquid Amies media.

The BioFire FilmArray Respiratory Panel 2 (RP2) is a multiplex in vitro molecular diagnostic test for the simultaneous and rapid detection of 21 pathogens, including 4 common human coronavirus strains, directly from nasopharyngeal swab (NPS) samples.

Stored residual serum and plasma samples submitted to the University of Chicago Medicine Clinical Laboratories for routine testing were recovered for this evaluation.

Percent agreement was determined and the hybrid Wilson/Brown method was used to calculate 95% confidence intervals of proportions (95% CI). All statistical analyses were performed using GraphPad Prism version 8.4.1.

## Results

Fifty-seven samples were tested from patients thought to be negative for exposure to SARS-CoV-2. Forty-one of the samples were from ambulatory patients at the University of Chicago with negative results for SARS-CoV-2 by PCR. The remaining 16 samples were collected in early 2019, prior to the current pandemic, and stored at -20°C. Twenty-eight of the 57 samples were from patients who had tested positive by the BioFire FilmArray RP2 respiratory viral panel for common coronavirus strains (6 samples positive for HKU1, 10 positive for NL63, 9 positive for OC43, 2 positive for 229E, and one positive for both OC43 and 229E).

Of these 57 samples, 53 tested negative and 4 tested positive for IgA (93.0% agreement, 95% CI: 83.3-97.2) while 56 tested negative and 1 tested positive for IgG (98.2% agreement, 95% CI: 90.7-99.9).

The sample that was positive for IgG was also positive for IgA. Chart review revealed an episode 4 weeks prior of a fever of 103.5°F and diarrhea, without cough. This clinical picture that could be consistent with COVID-19 disease. This positive serological result and negative PCR could represent exposure to SARS-CoV-2 with clearance of the virus.

Samples from sixty-seven unique specimens from 49 patients with PCR-positive SARS-CoV-2 were tested. Of these samples, 55 tested positive and 12 tested negative for IgA (82.1% agreement, 95% CI: 71.3-89.4) while 40 tested positive and 27 tested negative for IgG (59.7% agreement, 95% CI: 47.7-70.6). Six borderline positives for IgA and 2 borderline positives for IgG were included in the positive results.

These 67 samples from PCR positive patients were collected 0 to 45 days after PCR testing. The 12 IgA-negative samples and the 27 IgG-negative samples were collected within 7 days of PCR testing. Since antibody development occurs after viremia and requires time to reach a detectable concentration, it is likely that these samples were collected too early in the course of disease to expect antibody production.

The 67 serological samples from SARS-CoV-2 PCR-positive patients were stratified by the number of days following the positive PCR result. For samples collected ≥3 days after positive PCR, 29 of 30 (96.7%, 95% CI: 83.3-99.8) were positive for IgA (see chart 1); all 21 samples collected after one week were IgA positive. For samples collected ≥4 days after positive PCR, 28 of 28 (100%, 95% CI: 87.9-100) were positive for IgG (see chart 2).

**Chart 1.**
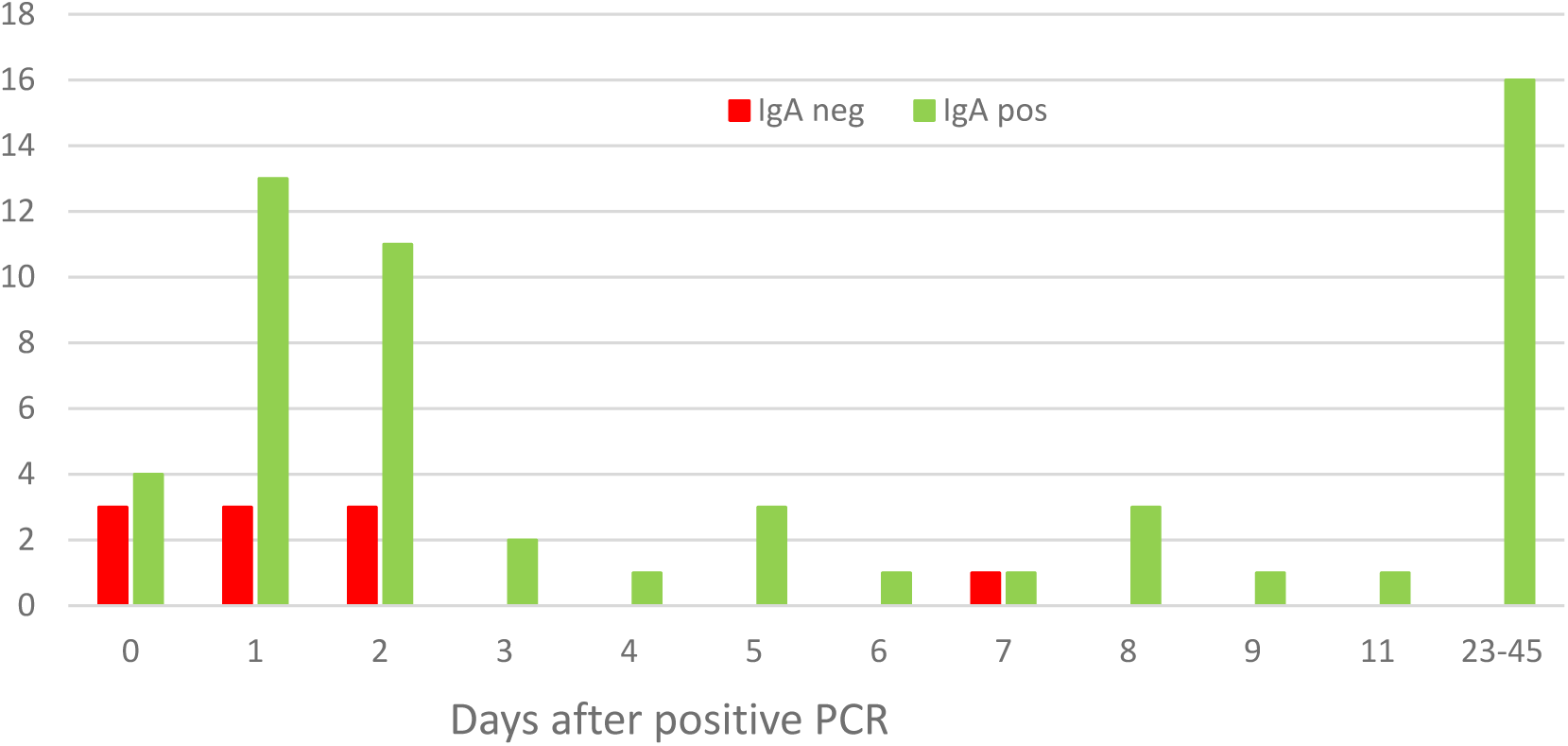
Timeline of IgA Results on PCR Positive Patients

**Chart 2.**
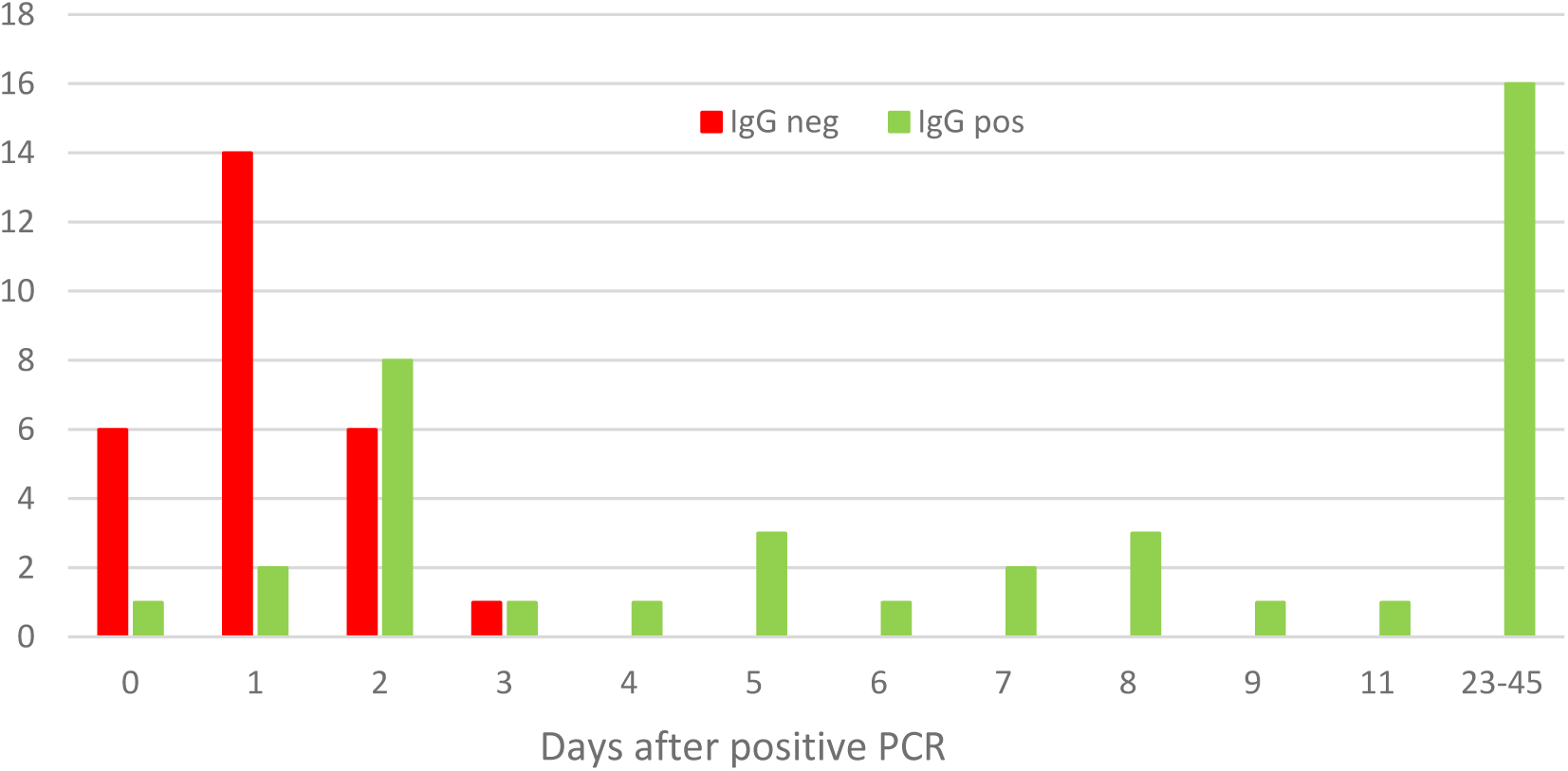
Timeline of IgG Results from PCR Positive Patients

## Cross-reactivity

A total of 28 unique samples that tested positive by BioFire FilmArray RP2 panel for common human coronaviruses including types 229E, NL63, HKU1 and OC43 were tested for cross reactivity. All 28 tested negative for both IgA and IgG antibodies.

## Discussion

A positive or negative test for the SARS-CoV2 antibody is difficult to interpret at this time. It is not yet known if antibodies serve as an indication of the presence or absence of protective or sustained immunity. Negative SARS-CoV-2 antibody results do not rule out SARS-CoV-2 infection, particularly in those who have been in contact with the virus. Results from antibody testing should not be used as the sole basis to diagnose or exclude SARS-CoV-2 infection or to inform infection status. Our evaluation of the EUROIMMUN assay did not demonstrate any cross reaction with samples from patients positive for common human coronaviruses.

## Conclusion

The EUROIMMUN Anti-SARS-CoV-2 ELISA Assay demonstrated excellent sensitivity for detection of IgA and IgG antibodies from samples collected >2 days and >3 days, respectively, after COVID-19 diagnosis by PCR. The EUROIMMUN Anti-SARS-CoV-2 ELISA Assay demonstrated excellent specificity and did not demonstrate cross reaction with common human coronaviruses, including types HKU1, NL63, CV229E, and OC43.

Limitations of this study include a small sample size and a lack of specimens collected more than 45 days days following a positive PCR.

## Acknowledgements

The authors thank Caroline Guenette and Caroline Hokl from Occupational Medicine at the University of Chicago.

## References

1. Lu R, Zhao X, Li J, Niu P, Yang B, Wu H, Wang W, Song H, Huang B, Zhu N, Bi Y, Ma X, Zhan F, Wang L, Hu T, Zhou H, Hu Z, Zhou W, Zhao L, Chen J, Meng Y, Wang J, Lin Y, Yuan J, Xie Z, Ma J, Liu WJ, Wang D, Xu W, Holmes EC, Gao GF, Wu G, Chen W, Shi W, Tan W. 2020. Genomic characterisation and epidemiology of 2019 novel coronavirus: implications for virus origins and receptor binding. Lancet 395:565–574. doi:10.1016/S0140-6736(20)30251-8.

2. Huang AT, Garcia-Carreras B, Hitchings MDT, Yang B, Katzelnick L, Rattigan SM, Borgert B, Moreno C, Solomon BD, Rodriguez-Barraquer I, Lessler J, Salje H, Burke DS, Wesolowski A, Cummings DAT. 2020. A systematic review of antibody mediated immunity to coronaviruses: antibody kinetics, correlates of protection, and association of antibody responses with severity of disease. medRxiv 04.14.20065771; doi: https://doi.org/10.1101/2020.04.14.20065771

3. Hsueh PR, Huang LM, Chen PJ, Kao CL, Yang PC. 2004. Chronological evolution of IgM, IgA, IgG and neutralisation antibodies after infection with SARS-associated coronavirus. Clin. Microbiol. Infect. 10:1062–1066.

4. Lassaunière R, Frische A, Harboe ZB, Nielsen ACY, Fomsgaard A, Krogfelt KA, Jørgensen CS. 2020. Evaluation of nine commercial SARS-CoV-2 immunoassays. medRxiv 04.09.20056325; https://doi.org/10.1101/2020.04.09.20056325

5. Okba NMA, Müller MA, Li W, Wang C, GeurtsvanKessel CH, Corman VM, Lamer MM, Sikkema RS, de Bruin E, Chandler FD, Yazdanpanah Y, Le Hingrat Q, Descamps D, Houhou-Fidouh N, Reusken CBEM, Bosch BJ, Drosten C, Koopmans MPG, Haagmans BL. 2020. Severe acute respiratory syndrome coronavirus 2-specific antibody responses in coronavirus disease 2019 patients. Emerg Infect Dis. Apr 8;26(7). doi: 10.3201/eid2607.200841

6. EUROIMMUN. 2020. Anti-SARS-CoV-2 ELISA IgG, Package insert. EI_2606G_A_US_C02.docx Version: 2020-05-04.

7. EUROIMMUN. 2020. Anti-SARS-CoV-2 ELISA IgA, Package insert.EI_2606A_A_US_C01.docx Version: 2020-03-24.

